# Complete Genome Sequences and Characteristics of Seven Novel Mycobacteriophages

**DOI:** 10.1101/2023.04.18.537415

**Authors:** Skylar M Weiss, Kezia K Happy, Faith W Baliraine, Abigail K Beach, Sean M Brobston, Claire P Martinez, Kaitlyn J Menard, Savannah M Orton, Angela L Salazar, Gregory D Frederick, Frederick N Baliraine

**Affiliations:** Department of Biology & Kinesiology, LeTourneau University, Longview, Texas, USA

## Abstract

Full genome sequences of seven mycobacteriophages isolated from environmental soil samples are presented. These bacteriophages, with their respective cluster or subclusters, are Duplo (A2), Dynamo (P1), Gilberta (A11), MaCh (A11), Nikao (K1), Phloss (N), and Skinny (M1). All were temperate Siphoviridae, with genome sizes ranging from 43,107–82,071 bp.

## ANNOUNCEMENT

Bacteriophages (aka <phages=) are viruses that exclusively infect bacteria, exhibiting obligate intracellular pathogenicity and a limited host range (1, 2). Bacteriophages are the most numerous entities in the biosphere, with an estimated population in excess of 10^31^ particles (3). The bacteria-bacteriophage relationship exhibits constant bidirectional selective pressure, with bacteria evolving to resist viral infection and bacteriophages coevolving to maintain replicative ability within their hosts (4, 5). The prevalence of antibiotic-resistant bacterial infections has propelled a resurgent interest in phage therapy (6–10). Here, we report on seven novel lysogenic bacteriophages.

All bacteriophages were isolated from environmental soil samples collected from various locations in East Texas during 2020-2021, using standard methods (5). In short, the soil samples were washed in Middlebrook 7H9 medium prior to centrifugation and supernatant filtration (0.22-μm pore size). The filtrates were subsequently inoculated with *Mycobacterium smegmatis mc*^*2*^*l55* and incubated at 25°C for 3.5 days with shaking. Filtered samples of each culture were plated with *M. smegmatis* in 7H9 top agar. After purification through three rounds of plating at 37°C for 48 hours, the observed plaque morphologies ranged from clear to turbid amongst the various phages (Table 1). Negative-stain transmission electron microscopy showed all these bacteriophages to exhibit a Siphoviridae morphotype with isometric capsids (diameter ~51.0 – 72.0 nm) and noncontractile tails (length ~124.0 – 281.0 nm; Fig. 1 & Table 1), measured using ImageJ (11–13).

**TABLE 1.**
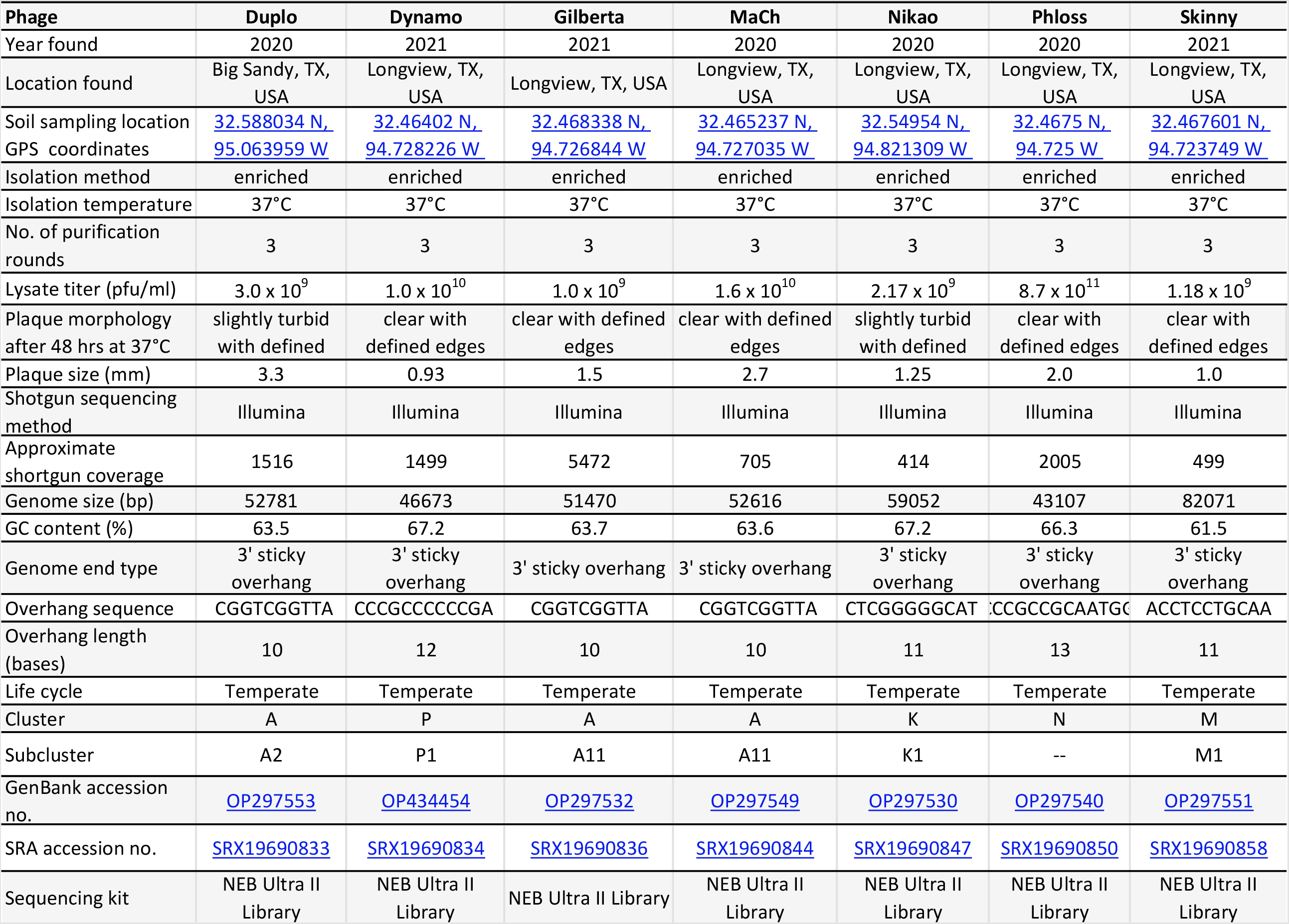
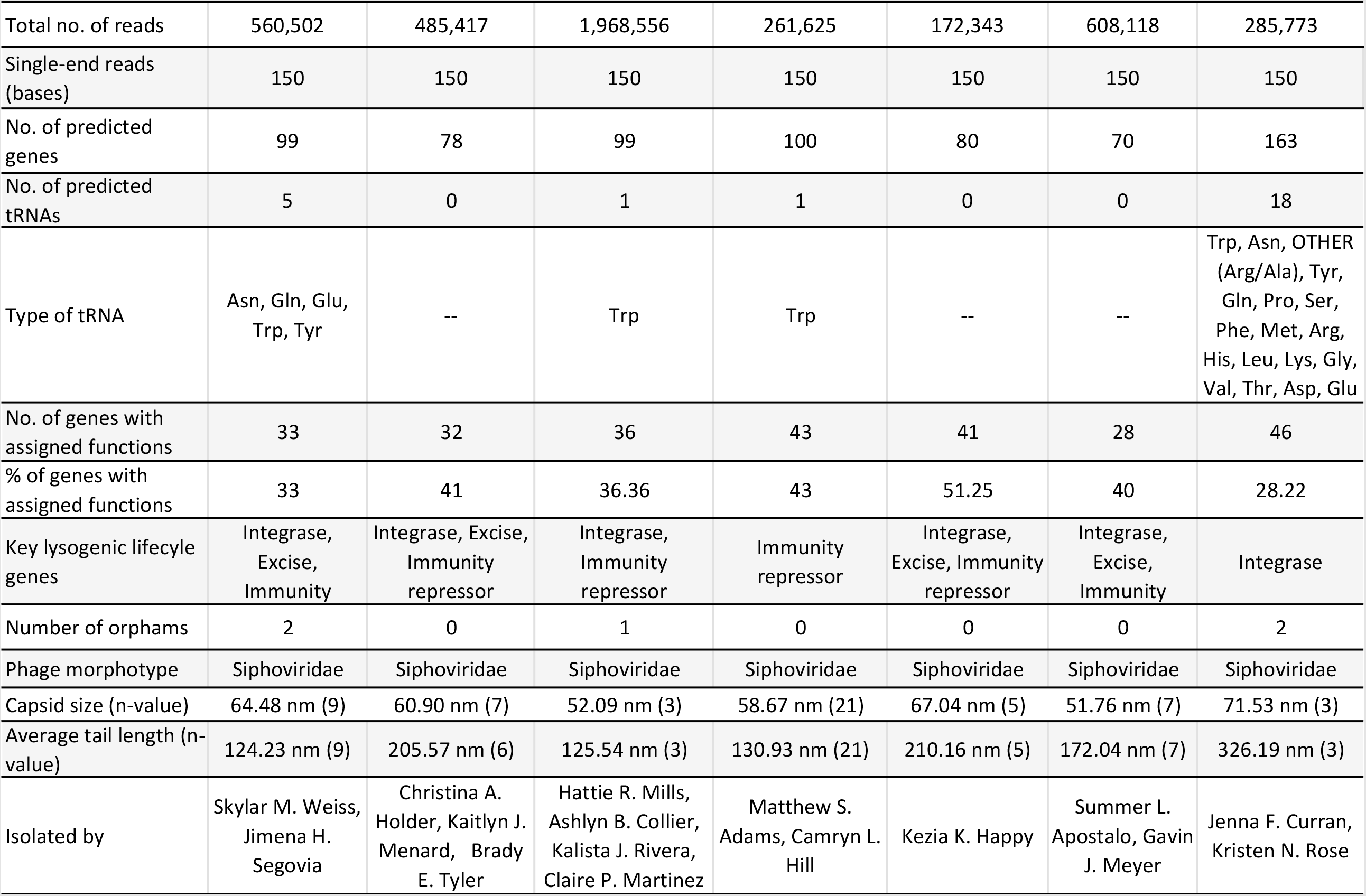
Properties of Seven Mycobacteriophages Isolated in East Texas in 2020 and 2021

**FIG 1.**
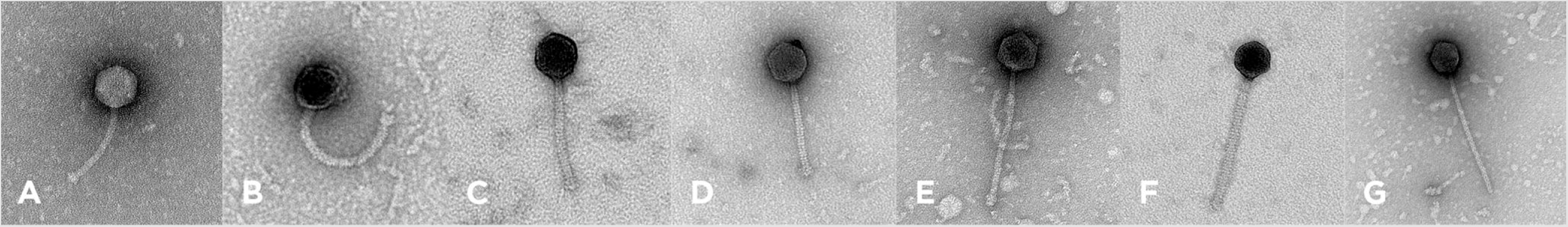
Transmission electron micrographs of the seven bacteriophages. (A) Duplo, (B) Dynamo, (C) Gilberta, (D) MaCh, (E) Nikao, (F) Phloss, and (G) Skinny. Capsid sizes and tail lengths are provided in Table 1. Bacteriophage particles were added to 300-mesh carbon–formvar-coated copper grids (Ted Pella Inc., Redding, CA), stained with 1% (wt/vol) uranyl acetate, and imaged at the University of Arkansas for Medical Sciences’ Digital Microscopy Laboratory.

Genomic DNA was extracted from lysates of varying titers (Table 1) using the Promega Wizard DNA cleanup kit. Preparation for sequencing using Illumina MiSeq (v3 reagents) utilized the NEB Ultra II Library kit. Assembly and verification of untrimmed reads was performed using Newbler v2.9 (14) and Consed v29 (15, 16). Sequencing revealed genomes ranging in length from 43,107 bp in Phloss to 82,071 bp in Skinny (Table 1). All had 3’ sticky overhangs (10-13 bp long) with an average G+C of 64.7% (range 61.5–67.2%) comparable to the 67.4% G+C content of their isolation host *Mycobacterium smegmatis* mc^2^155 (17). The seven phages were assigned to subclusters A2, A11, K1, M1, P1, and cluster N (Table 1) based on ≥35% gene content similarity (GCS) to other phages, using the GCS tool in PhagesDB (18, 19).

Genome annotation was accomplished using <u>DNAMaster</u> v5.23.6, <u>Starterator</u>, Phamerator (20), BLASTp in NCBI and PhagesDB (21, 22), GenMark v2.5p (23), HHpred (24, 25), Glimmer v3.02 (26), TMHMM v.2.0 (27), SOSUI v1.11 (28), tRNAscan-SE v2.0 (29, 30), and ARAGORN v1.2.41 (31). Default settings were utilized within all programs (32). An average number of 98.0 putative protein-coding genes (range 70-163) and 3.6 tRNAs (range 0-18) were predicted (Table 1). Functions could only be assigned to 28-51% of the putative genes across the phages (Table 1). All phages had at least one of the three key genes associated with a lysogenic lifecycle. Duplo, Dynamo, Nikao, and Phloss had the integrase, excise, and immunity repressor, Gilberta had both the integrase and immunity repressor, while Skinny and MaCh had only the integrase and the immunity repressor genes, respectively (Table 1).

## Data and lysate availability

Raw reads of all seven reported mycobacteriophages are available in the Sequence Read Archive (SRA) database, and their complete genome sequences are available in GenBank database. Their SRA and GenBank accession numbers with their respective Uniform Resource Locators (URLs) are provided in Table 1. High titer lysates of the phages are archived at the University of Pittsburgh Bacteriophage Institute.

## ACKNOWLEDGEMENTS

We thank Graham Hatfull for his enduring commitment to and leadership of the Science Education Alliance-Phage Hunters Advancing Genomics and Evolutionary Science (SEA-PHAGES) program, Debbie Jacobs-Sera, Viknesh Sivanathan for their expertise and support, and Daniel Russell and Rebecca Garlena of the University of Pittsburgh Bacteriophage Institute for sequencing and assembling the phage genomes. We appreciate Brianna Mack and Jonathan Matthews for preparing the laboratory materials used. We thank Matthew Adams, Summer Apostalo, Camryn Hill, Ashlyn Collier, Jenna Curran, Gavin Meyer, Christina Holder, Kaitlyn Menard, Hattie Mills, Jimena Segovia, Brady Tyler, Kalista Rivera, and Kristen Rose who helped isolate the phages. We thank Jeff Kamykowski of the University of Arkansas for Medical Sciences Digital Microscopy Laboratory for capturing the transmission electron micrographs for these phages. Many thanks to the Howard Hughes Medical Institute for their support of the SEA-PHAGES program, and to LeTourneau University School of Arts & Sciences for supporting and facilitating this undergraduate bacteriophage research.

